# The Endosomal pH Regulator NHE9 is a Driver of Stemness in Glioblastoma

**DOI:** 10.1101/2021.10.07.463493

**Authors:** Myungjun Ko, Monish R. Makena, Paula Schiapparelli, Paola Suarez-Meade, Allatah X. Mekile, Bachchu Lal, Hernando Lopez-Bertoni, John Laterra, Alfredo Quiñones-Hinojosa, Rajini Rao

## Abstract

A small population of self-renewing stem cells initiate tumors and maintain therapeutic resistance in glioblastoma. Given the limited treatment options and dismal prognosis for this disease there is urgent need to identify drivers of stem cells that could be druggable targets. Previous work showed that the endosomal pH regulator NHE9 is upregulated in glioblastoma and correlates with worse survival prognosis. Here, we probed for aberrant signaling pathways in patient-derived glioblastoma cells and found that NHE9 increases cell surface expression and phosphorylation of multiple receptor tyrosine kinases by promoting their escape from lysosomal degradation. Downstream of NHE9-mediated receptor activation, oncogenic signaling pathways converged on the JAK2-STAT3 transduction axis to induce pluripotency genes Oct4 and Nanog and suppress markers of glial differentiation. We used both genetic and chemical approaches to query the role of endosomal pH in glioblastoma phenotypes. Loss-of-function mutations in NHE9 that failed to alkalinize endosomal lumen did not increase self-renewal capacity of gliomaspheres *in vitro*. However, monensin, a chemical mimetic of Na^+^/H^+^ exchanger activity, and the H^+^ pump inhibitor bafilomycin bypassed NHE9 to directly alkalinize the endosomal lumen resulting in stabilization of receptor tyrosine kinases and induction of Oct4 and Nanog. Using orthotopic models of primary glioblastoma cells we found that NHE9 increased tumor initiation *in vivo*. We propose that NHE9 initiates inside-out signaling from the endosomal lumen, distinct from the established effects of cytoplasmic and extracellular pH on tumorigenesis. Endosomal pH may be an attractive therapeutic target that diminishes stemness in glioblastoma, agnostic of specific receptor subtype.

**Significance:** A well-known hallmark of cancer is excessive acidification of tumor microenvironment, caused by upregulation of Na^+^/H^+^ exchanger activity on the cancer cell membrane. However, the role of organellar pH in tumor biology has not been established. This study identifies a mechanistic link between upregulation of the endosomal Na^+^/H^+^ exchanger NHE9 and stemness properties in glioblastoma, the most malignant and common brain tumor in adults. By increasing pH of the recycling endosome, NHE9 exerts a broad effect on post-translational stability and activation of multiple receptor tyrosine kinases, leading to increased stem cell-like properties of self-renewal and tumor initiation in glioblastoma models. Our findings suggest that targeting NHE9 or endosomal pH could be an effective strategy for receptor agnostic glioblastoma treatment.

## INTRODUCTION

Glioblastoma (GBM) is the most frequent type of brain tumor in adults and the most malignant form (grade IV) of glioma with average 3-year survival rate of less than 10% (1–3). The currently available “gold standard” for GBM patients is surgical resection of the tumor mass, followed by radiotherapy and concurrent chemotherapy with the alkylating agent temozolomide (TMZ)(4–6). Median survival with TMZ treatment and radiotherapy combined after surgical intervention has only increased marginally from 12 months to 14.6 months (3, 7). The problem is that there are no other chemotherapeutic drugs or targets that have been approved and effective for GBM patients. Thus, a global effort in the brain tumor field has been focused on finding novel therapeutic targets for this rapidly fatal brain tumor. Of central importance in these investigations are cancer stem cells, a subset of cancer cells with stem cell-like characteristics, that may be responsible for the resistance of GBM to conventional therapies and recurrence of the disease (8, 9). There is a high unmet need to find novel therapeutic targets that are drivers of cancer stem cells.

Aberrant signaling pathways and epigenetic changes are the main drivers of GBM tumor phenotype and resistance to therapy. Receptor tyrosine kinases (RTKs) are cell surface receptors for growth factors, hormones, cytokines and neurotrophic factors that regulate tumor cell proliferation, migration, differentiation and survival through downstream signaling pathways. Although expression and activation of RTKs is frequently aberrant in GBM, redundant signaling pathways and tumor heterogeneity in RTK expression make it extremely hard for RTK-targeted therapy to be effective (10, 11). Combination therapy - targeting multiple RTKs, has been suggested (12) but the plasticity of tumor cells and their stochastic response has prevented this approach from being effective and approved for patients. Furthermore, many independent groups have uncovered a small population of cells in GBM that have stem cell-like characteristics that confer treatment resistance and self-renewal (8, 13). Thus, a druggable target that controls both stemness characteristics and pan-receptor clearance in GBM cells could circumvent shortcomings in current therapy.

A perturbation in pH dynamics is a common pathophysiological hallmark of tumors (14–16). The accumulation of acidic byproducts of metabolism together with poor vascularization and accompanying hypoxia demands net acid extrusion from the cancer cell, which drives tumor microenvironment to pH 6.5 or lower. One well-studied mechanism underlying these pH changes is activation of the plasma membrane Na^+^/H^+^ exchanger NHE1, which plays a central role in tumor cell migration and invasion (17, 18). In contrast, several poorly-understood intracellular NHE isoforms (19) that regulate the lumenal pH of endosomes, Golgi and *trans*-Golgi network are emerging as potential drivers of tumorigenesis and drug resistance, and offer exciting new mechanistic insights that could be therapeutically exploited (20–22).

One such intracellular isoform is NHE9, which localizes to recycling endosomes and is widely expressed in the brain (23, 24). Even small changes in Na^+^/H^+^ exchange activity can cause large shifts in pH within the limited confines of the endosomal lumen (20, 24). NHE9 has been implicated in multiple malignancies, including glioblastoma (20), prostate cancer (25), colorectal cancer (26–28), esophageal squamous carcinoma (29–32), oral cancer (33), and ovarian cancer (34). However, the underlying mechanistic basis for these observations is largely unclear.

In this study we sought to broaden our initial observation that NHE9 increases membrane persistence of the epidermal growth factor receptor EGFR in glioblastoma cells (20). Our new findings suggest that NHE9 may have a more global effect on receptor stabilization that is agnostic of GBM subtype. We found JAK2/STAT3 serves as a common signal transduction pathway downstream of NHE9-mediated receptor tyrosine kinase activation. STAT3 is a critical signaling molecule (35–37) that controls a small population of self-renewing stem cells that occupy the apex of the tumor cell hierarchy and drive initiation (38, 39), growth (40), and therapeutic resistance (8, 41) in glioblastoma. These observations led us to define the intermediate steps linking NHE9 to self-renewal capacity in gliomaspheres *in vitro*, and tumor initiation *in vivo.* Both genetic and pharmacological tools point toward a causal role for endosomal pH in glioblastoma stem cell-like properties. In summary, we report that “inside-out” signaling from the endosomal lumen can drive stemness, which could explain poor survival prognosis glioblastoma patients with elevated NHE9 expression. We suggest that inhibition of NHE9 will attenuate oncogenic signaling in a receptor-agnostic manner, opening potential avenues for cancer therapy.

## RESULTS

### NHE9 stabilizes multiple receptor tyrosine kinases

Following endocytosis, internalized ligand-receptor complexes may be recycled back to the cell surface, delivered to the lysosome for degradation, or dissociate to undergo divergent fates. Signaling molecules like growth factors are typically degraded along with their cognate receptors as a critical step in down-regulation of cellular response. Multiple steps in this process are exquisitely dependent on the acidic pH of the endosomal lumen, including dissociation of ligand-receptor complex, cargo sorting decisions, and vesicle docking and fusion. Previously, we showed that NHE9 increased membrane persistence of EGFR in glioblastoma cells (20). Here, we hypothesize that dysregulation of pH in the recycling endosome in glioblastoma cells, due to increased Na^+^/H^+^ exchange activity globally elevates surface expression of receptor tyrosine kinases (RTK), leading to prolonged oncogenic signaling (Fig. 1A).

**Figure 1:**
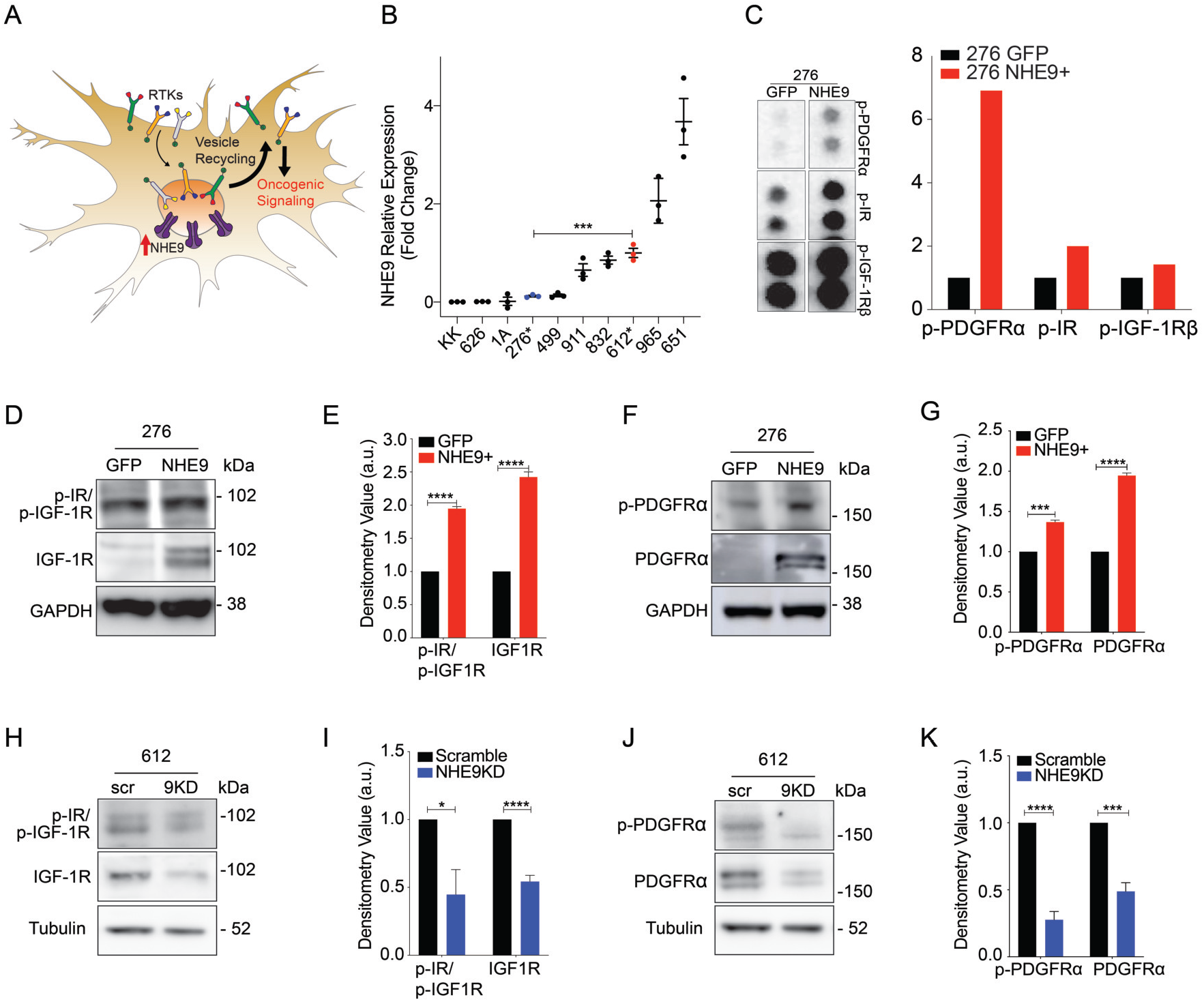
NHE9 increases activation and expression of receptor tyrosine kinases. (A) Schematic showing the hypothesis that increased endosomal NHE9 facilitates recycling of the receptor tyrosine kinases (RTKs) to prolong downstream oncogenic signaling. (B) Relative mRNA expression of NHE9 measured by qPCR for ten GBM cell lines established from samples obtained from Johns Hopkins neurosurgery operating room. GBM 276 and GBM 625 were chosen to represent low and high expressors, without being outliers. (C) Portion of phospho-RTK dot blot (left) probed with lysates from GBM 276 ectopically expressing GFP or NHE9-GFP. Densitometry quantification (in arbitrary units) of p-PDGFRα, p-IGF-1Rβ, and p-IR from the dot blot (right). (D, F) Representative Western blot showing p-IGF-1R, p-IR, and p-PDGFRα (top row) and IGF-1R, IR, and PDGFRα (bottom row) for GBM 276 GFP or NHE9-GFP and densitometry quantification from biological triplicates (E, G). (H, J) Representative Western blot showing p-IGF-1R, p-IR, and p-PDGFRα (top row) and IGF-1R, IR, and PDGFRα (bottom row) for GBM 612 scramble or shNHE9 and densitometry quantification from biological triplicates (I, K).

To test this hypothesis, we first screened for NHE9 expression a diverse panel of 10 patient-derived glioblastoma cell lines, established from samples obtained from the operative room at the Johns Hopkins Hospital and propagated in serum-free medium (Fig. 1B). We selected the GBM cell lines, GBM 276 and GBM 612, as representing low and high expression of endogenous NHE9, without being outliers. GBM 276 and GBM 612 represent distinct glioblastoma subtypes-proneural and mesenchymal, respectively (20). We further generated isogenic matched pairs of GBM cells representing loss or gain of NHE9 function using lentiviral shRNA for NHE9 knockdown (KD) in GBM 612 and transgenic expression of NHE9 tagged with GFP in GBM 276 (Fig. S1A-C). Previously, we showed that endosomal pH correlated with NHE9 expression levels in GBM cells (20). Specifically, endosomal pH was lower in GBM 276 (5.57 ± 0.13) compared to GBM 612 (6.88 ± 0.13), consistent with a role for NHE9 in limiting lumen acidification. To ensure that transgene expression of NHE9-GFP physiologically mimics the endogenous high levels of NHE9 in GBM, we demonstrated that endosomal localization of NHE9-GFP in GBM 276 was accompanied by increase in lumen pH to 6.61 ± 0.21, similar to GBM 612. Notably, no change in cytoplasmic pH was observed in response to manipulation of NHE9 expression (20). Thus, these cells are well-suited for mechanistic studies of NHE9 in glioblastoma.

Transgene expression of NHE9 in GBM 276 resulted in the activation of multiple receptors detected on a phospho-RTK dot blot overlaid with GBM cell lysate (Fig. S1D). The most prominent of these include insulin-like growth factor receptor 1 (IGF-1R), insulin receptor (IR), and platelet-derived growth factor receptor alpha (PDGFRα), (Fig. 1C). Western blot analysis of lysates from GBM 276 cells confirmed that gain of function in NHE9 increased phosphorylation in these receptors (Fig. 1D-G, top panels). Conversely, these receptors showed diminished phosphorylation in response to NHE9 knockdown in GBM 612 (Fig. 1H-K, top panels). Strikingly, gain or loss of NHE9 also caused corresponding increase or decrease in total levels RTK proteins (Fig. 1D-K, bottom panels). Notably, transcript levels of these receptors remained unchanged by the expression of NHE9, as determined by qPCR (Fig. S1 E-G) pointing to post-translational mechanisms underlying the differences in receptor levels. Therefore, we hypothesized that NHE9 altered receptor stability and turnover.

To probe cell surface expression of RTKs, we used flow cytometry analysis of non-permeabilized GBM 276 cells labeled with antibody against external receptor epitopes. We observed significant right shift of IGF-1R (Fig. 2A) or PDGFRα (Fig.2B) indicating increased surface expression in response to ectopic expression of NHE9, consistent with our model (Fig. 1A). Immunofluorescence imaging revealed co-localization of IGF-1R with NHE9-GFP in GBM 276 cells (Fig. 2C), reinforcing our hypothesis. Next, we treated cells with MG132 and bafilomycin, to block proteasomal and lysosomal degradation, respectively. In control GBM 612 cells, knockdown of NHE9 decreased levels of IGF-1R and PDGFRα, similar to results shown in Fig. 1. Whereas MG132 had no effect on RTK levels, treatment with bafilomycin was effective in protecting IGF-1R and PDGFRα protein levels in NHE9 KD cells (Fig. 2D). Taken together with our earlier findings on EGFR persistence in GBM (20), these observations suggest that NHE9 promotes RTK escape from lysosomal degradation.

**Figure 2:**
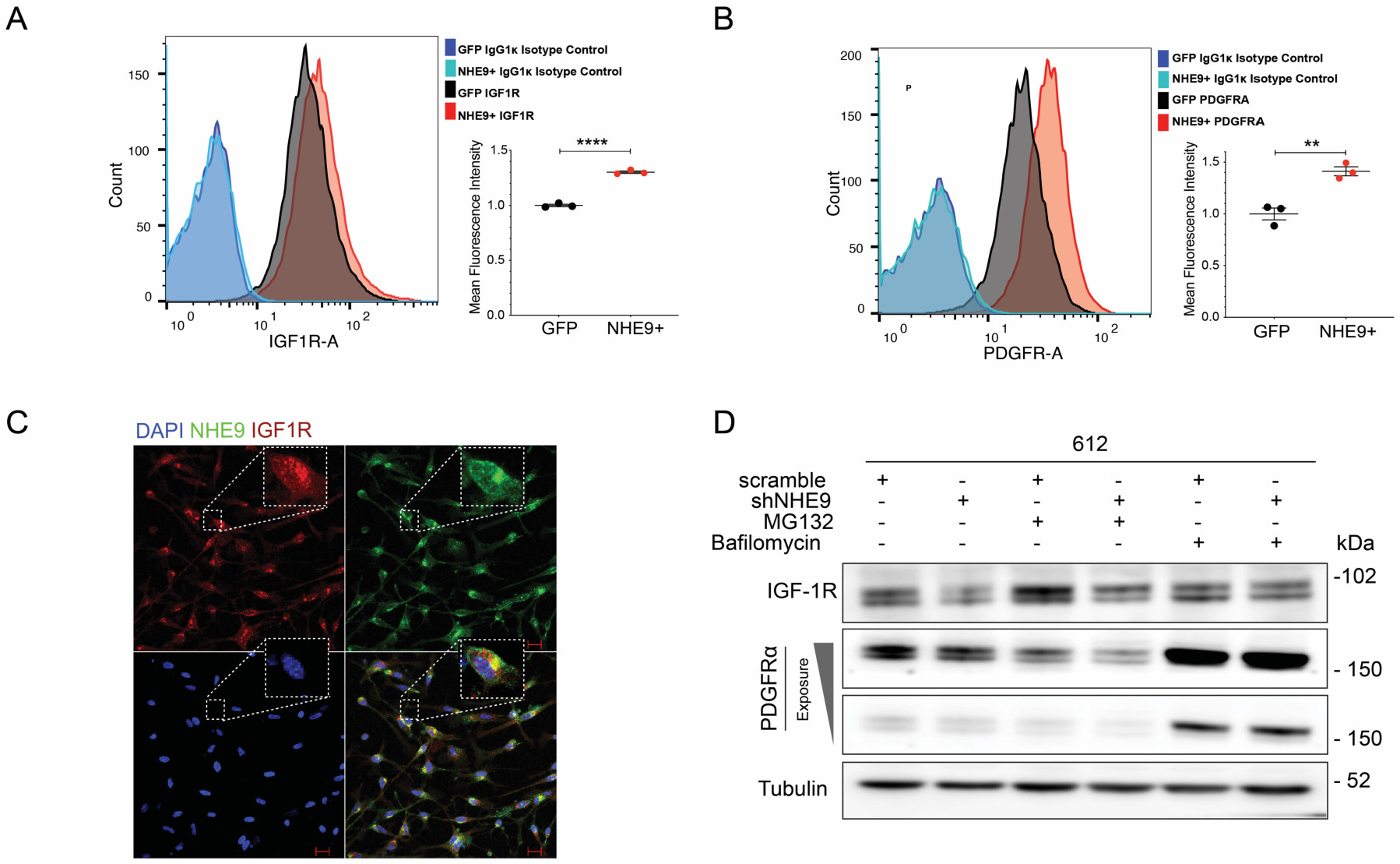
NHE9 increases surface expression and decreases lysosomal degradation of receptor tyrosine kinases. (A-B) Representative flow cytometry analysis of non-permeabilized GBM 276 cells expressing GFP or NHE9-GFP, treated with monoclonal antibodies for IGF-1R (A) and PDGFRα (B) in comparison with IgG1 kappa isotope control. Mean fluorescence intensity of peak areas are from biological triplicates. (C) Immunofluorescence images of GBM 276 cells showing IGF-1R (Red), NHE9-GFP (Green) and DAPI (blue). Merge of all three labels shows co-localization of IGF-1R and NHE9 (yellow). Scale bar: 10μm. (D) Western blotting analysis of IGF-1R and PDGFRα in GBM 612 scramble or NHE9KD cells treated with 25 nM MG132 or 25 nM bafilomycin for 6 h. Note that only bafilomycin diminishes the effect of shNHE9 on receptor levels.

### NHE9 induces phosphorylation of STAT3, a key prognostic marker in glioblastoma

To identify potential links between NHE9 and oncogenic pathways functioning downstream of RTK activation, we screened the TCGA pan-cancer studies dataset for signaling protein expression against NHE9 transcript levels. STAT3 is widely recognized as a master regulator and driver of transforming events leading to GBM, and has been studied as a prognostic marker and therapeutic target in glioblastoma (37, 42, 43) and other cancers (36, 44). We noted that p-STAT3 (Y705) had the highest correlation with NHE9 expression (Fig. 3A). The same analysis was performed in glioma patients for pSTAT3 (Y705), again revealing significant correlation with NHE9 (Fig. 3B). Co-expression analysis in glioblastoma patients revealed a modest, albeit significant positive correlation between NHE9 mRNA and the STAT signaling axis, including JAK2 and STAT3 (Pearson’s coefficient: 0.262 and 0.163 respectively) (Fig. S2A). These observations identified STAT3 signaling as a potential downstream mediator of NHE9 in glioblastoma (Fig. 3C).

**Figure 3:**
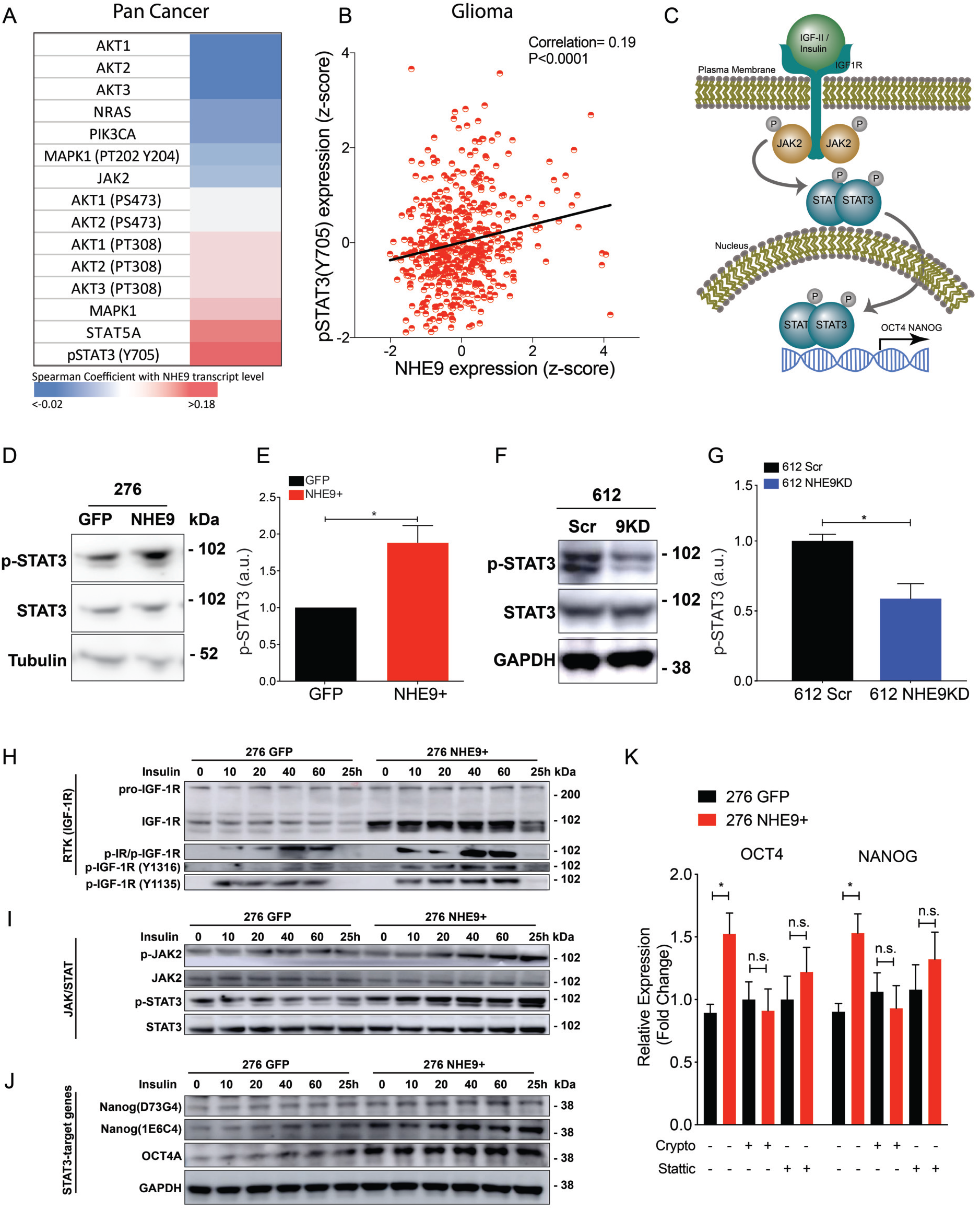
NHE9 activates the STAT3 signaling axis. (A) Heatmap of Spearman coefficient correlating protein expression levels of various oncogenic signaling proteins and NHE9 transcript in the TCGA pan-cancer dataset. (B) Correlative analysis between p-STAT3 (Y705) protein level and NHE9 transcript in the TCGA glioma patient cohort. (Correlation coefficient: 0.19; \*\*\*\**p-*value < 0.0001) (C) Schematic of the IGF-1R-JAK2-STAT3 signaling axis. (D) Representative Western blot of p-STAT3 and STAT3, and densitometry quantification (E). (F) Representative Western blot GBM 612 scramble or NHE9KD probing for p-STAT3 and STAT3 and its densitometry quantification (G). (H-J) Western blotting analysis with time course for the indicated time points in minutes, and 24 hr post ligand treatment with insulin in GBM 276 GFP or NHE9-GFP as described in Methods. Blots were probed for (H) pro-IGF-1R, IGF-1R, p-IR/IGF-1R, p-IGF-1R (Y1216), p-IGF-1R (Y1135), (I) p-JAK2, JAK2, p-STAT3, STAT3, (J) Nanog (D73G4), Nanog (1E6C4), OCT4A, and GAPDH. (K) qPCR analysis for Oct4 and Nanog transcript in GBM 276 GFP or NHE9-GFP with or without STAT3 inhibitors, Cryptotanshinone (Crypto) or Stattic.

We tested this hypothesis by directly probing STAT3 phosphorylation in GBM 276 and 612 cells in response to NHE9 overexpression and knockdown, respectively. Western blotting of cell lysates revealed that ectopic expression of NHE9 significantly elevated p-STAT3 (pY705) (Fig. 3D-E) whereas attenuation of NHE9 expression led to corresponding decrease in STAT3 phosphorylation (Fig. 3F-G). These findings suggest a causal relationship between NHE9 expression levels and STAT3 signaling that warranted further analysis.

We used Western blotting to query the IGF-1R signaling cascade (Fig. 3C) in response to insulin (25 μg/ml) treatment of GBM 276 cells. Total IGF-1R expression was elevated in transgene expressing GBM 276 NHE9+ cells, compared to GFP-transfected control (276 GFP) at all time points after ligand treatment, with significant turnover of the receptor observed at 24 h (Figure 3H). The activated form of IGF-1R was observed by using three different antibodies that detect p-IR/p-IGF-1R (Y1150/1151 and Y1135/1136, respectively), p-IGF-1R (Y1316), and p-IGF-1R (Y1135), respectively. In each of the antibody blots, p-IGF-1R level in response to insulin addition was enhanced in NHE9+ cells but returned to non-stimulated levels by 24 h (Fig. 3H). Similarly, time-dependent increase of p-STAT3 and p-JAK2 was observed following insulin addition, with higher phosphorylation levels in GBM 276 NHE9+ cells (Fig. 3I). Remarkably, persistent phosphorylation at 24 h post-insulin addition was suggestive of prolonged downstream effects of RTK activation. Total levels of STAT3 and JAK2 remained unchanged in both cell lines, again pointing to a post-translational effect. Later in the time course, expression of two downstream targets of the JAK-STAT pathway, Oct4 and Nanog, was achieved in response to STAT3 elevation, including and up to 24 h after insulin treatment (Figure 3J). Target gene expression was significantly enhanced in NHE9+ cells, whereas the housekeeping protein GAPDH was similar under all conditions (Fig. 3J).

We used pharmacological blockers to validate the role of STAT3 signaling in the transcriptional output of Oct4 and Nanog downstream from NHE9. Inhibition of p-STAT3 with Stattic and Crypto (at 6.25 and 10 μM, respectively) successfully abrogated the effect of NHE9 on Oct4 and Nanog expression as evidenced by their mRNA (Fig. 3K) and protein levels (Fig. S2B). The efficacy of these inhibitors and their dose-dependent ability to decrease p-STAT3 levels in GBM 276 was confirmed (Fig S2C-E).

### NHE9 induces pluripotency genes and suppresses differentiation markers

Oct4 and Nanog, also known as pluripotency factors, are transcription factors highly expressed in embryonic stem cells (45), and capable of reprogramming cells of committed lineage back to their primitive, stem cell state (46–48) (Fig. 4A). These reprogramming factors are also highly expressed in malignancies such as glioblastoma, where they confer worse prognosis in patients by eliciting therapeutic resistance and disease recurrence (49, 50). Consistent with the results from Fig. 3, we observed significant elevation of both Oct4 and Nanog transcripts in GBM 276 NHE9+ cells (Fig. 4B) and conversely, significant downregulation in GBM 612 NHE9 KD cells generated independently with lentiviral mediated shRNA (Fig. 4C) or siNHE9 coated poly-β-amino ester (PBAE) nanoparticles (Fig. 4D). The effect on Oct4 and Nanog was selective as expression of Sox2, which is also linked to stemness, showed modest, or no significant change (Fig. 4B, C). Western blot analysis was used to confirm that protein levels of Oct4 and Nanog were elevated in response to transgene NHE9 expression in GBM 276 (Figure 4E - H), and reciprocally, decreased upon NHE9 knockdown in GBM 612 (Figure 4I - L).

**Figure 4:**
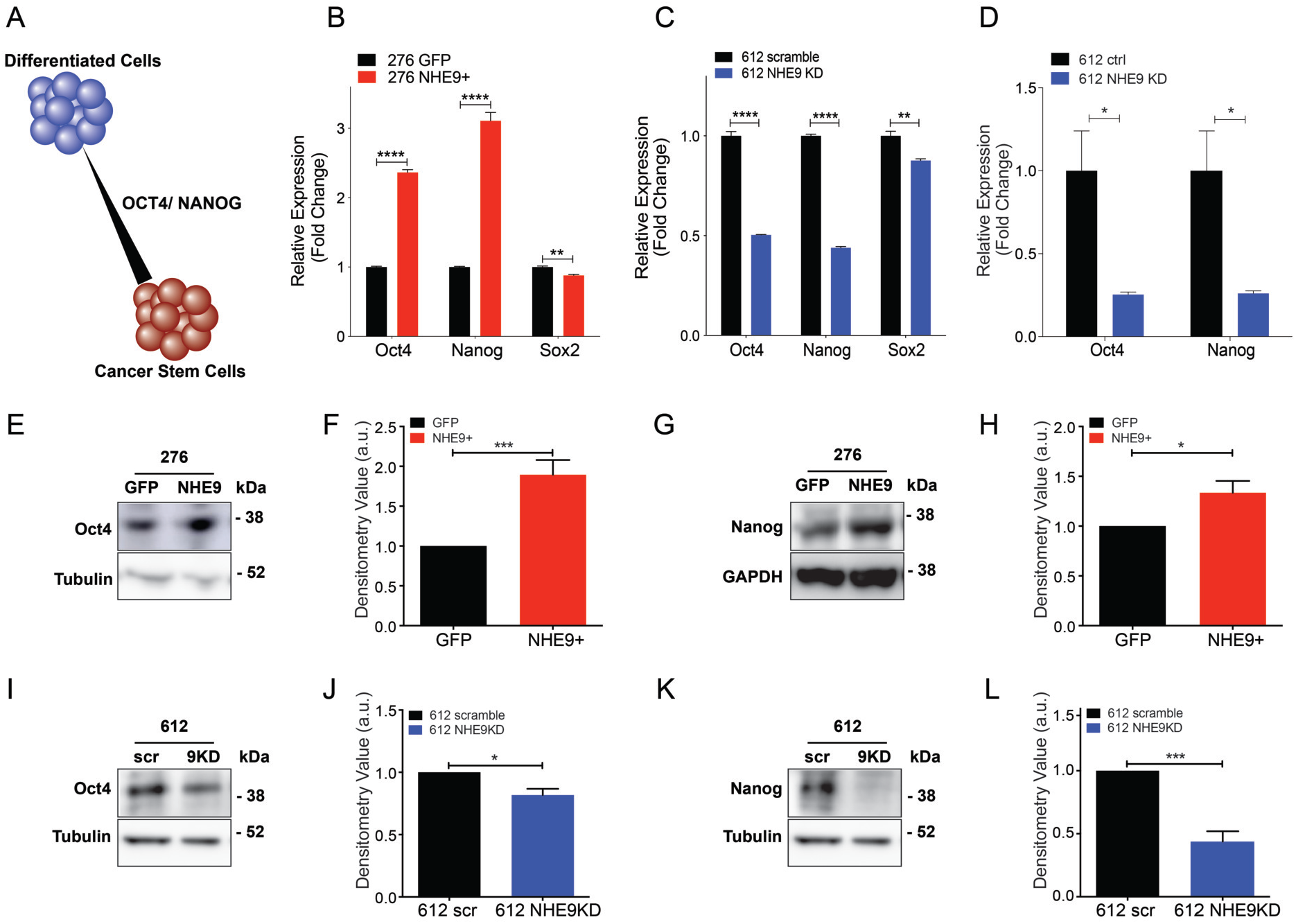
NHE9 increases stemness gene expression. (A) Schematic showing that Oct4 and Nanog shift distribution of cell state towards cancer stem cells (red), relative to differentiated cells (blue). (B-D) qPCR analysis of Oct4, Nanog, and Sox2 transcripts, as indicated. (B) GBM 276 GFP and NHE9-GFP (C) GBM 612 scramble and shNHE9 (NHE9 KD) (D) GBM 612 control siRNA or siNHE9-coated poly-β-amino ester (PBAE) nanoparticles (NHE9 KD). Averages of three biological replicates shown. Representative Western blots for (E) Oct4 and (G) Nanog in GBM 276 GFP or NHE9-GFP and (F, H) densitometry quantifications from three biological replicates. (I-L) Representative Western blots for (I) Oct4 and (L) Nanog in GBM 612 scramble or NHE9 KD and densitometry quantifications from three biological replicates (J, L).

Next, we asked whether NHE9 expression was linked to reciprocal changes in differentiation markers that identify astrocyte lineage in GBM cells (Fig. 5A). We noted that transcript levels of the astrocytic marker glial fibrillary acidic protein (GFAP) were significantly higher in GBM 276 compared to GBM 612, corresponding to low and high endogenous NHE9, respectively (Fig. 5B). Further evaluation of matched NHE9 +/− pairs showed that expression of lineage markers for oligodendrocytes (O4), neurons (Tuj1), and astrocytes (GFAP) was decreased in GBM 276 NHE9+ cells (Fig. 5C), whereas NHE9 knockdown in GBM 612 increased transcript of all three lineage markers (Fig. 5D). Protein analysis recapitulated the qPCR results: decreased GFAP protein levels in GBM 276 NHE9+ cells as seen on Western blots (Fig. 5E, F) and by confocal immunofluorescence microscopy (Fig. 5G, H). Conversely, NHE9 KD in GBM 612 increased GFAP level observed by Western blotting (Fig. 5I, J) and immunofluorescence (Fig. 5K, L). Together, these results consistently demonstrate that NHE9 elicits stemness markers and may shift glioblastoma to a cellular state closer to progenitor stem cells.

**Figure 5:**
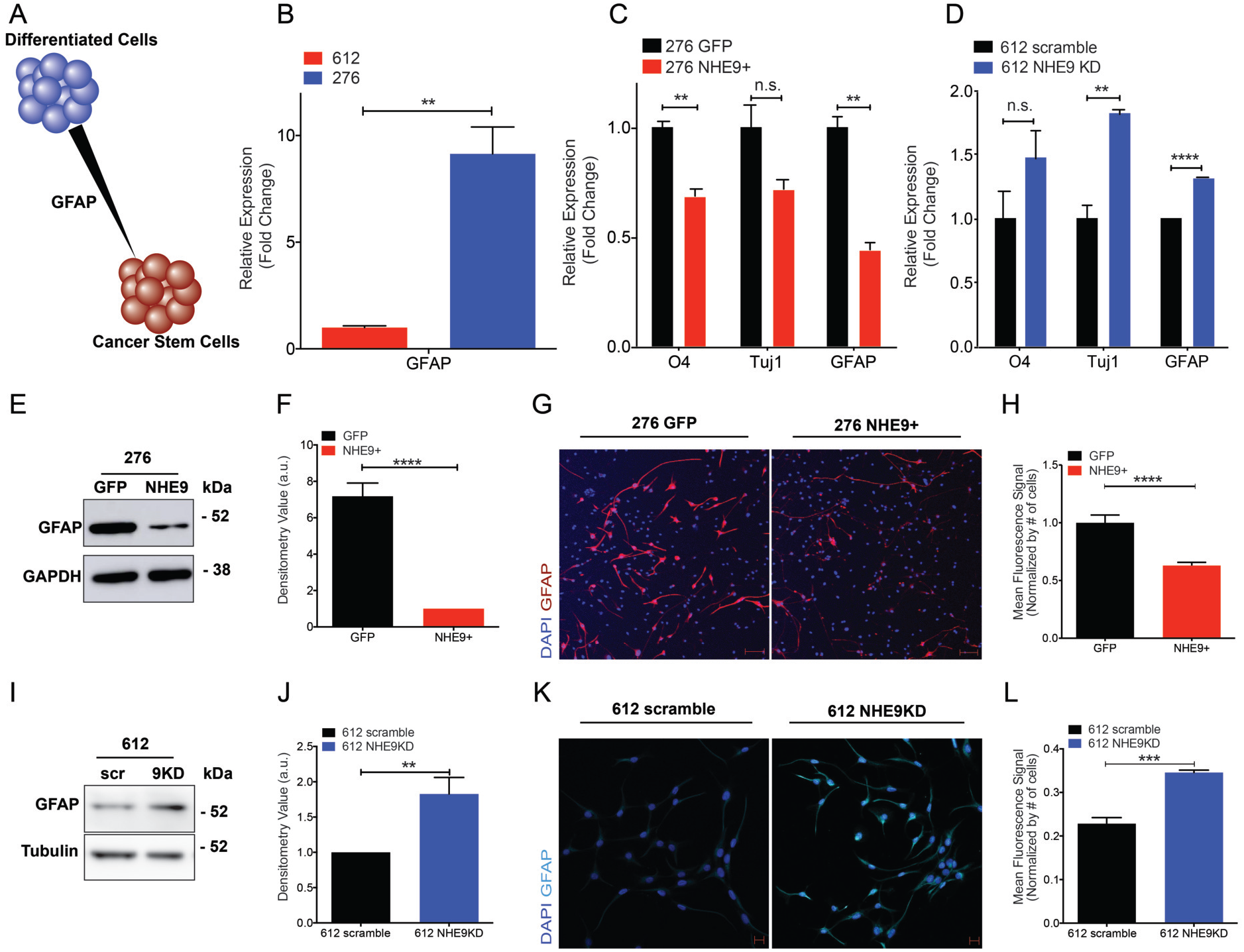
NHE9 decreases expression of differentiation markers. (A) Schematic showing that lineage markers such as GFAP shift cancer stem cells towards a differentiated state. (B-D) qPCR analysis for the indicated lineage markers, averaged from three biological replicates. (B) GBM 612 and GBM 275 (C) GBM 276 GFP or NHE9-GFP (D) GBM 612 scramble or NHE9 KD. (E) Representative Western blot for GFAP in GBM 276 GFP or NHE9-GFP and (F) densitometry quantification from three biological replicates. (G-H) Representative immunofluorescence image for (G) GFAP (red) and DAPI (blue) in GBM 276 GFP and NHE9-GFP and (H) fluorescence signal quantification. (I) Representative Western blot for GFAP in GBM 612 scramble or NHE9-GFP and (J) densitometry quantification from three biological replicates. (K-L) Representative immunofluorescence image for (K) GFAP (light blue) and DAPI (blue) in GBM 612 scramble or NHE9 KD and (L) mean fluorescence signal quantification.

### Endosomal alkalinization is required for glioblastoma stemness

We used both genetic and chemical approaches to manipulate endosomal pH (Fig. 6A) and evaluate stemness phenotypes. First, we considered whether ion transport by NHE9 was required for driving stemness characteristics. In addition to the membrane embedded transporter domain, Na^+^/H^+^ exchangers have a cytoplasmic tail (~150 aa), known to bind regulatory factors and scaffold signaling proteins that are implicated in tumorigenesis (21, 29, 51). Thus, the ability of NHE9 to drive stemness may be due to ion transport activity and/or signaling through the cytoplasmic tail. To begin to distinguish between these roles, we deployed previously characterized, autism-associated missense NHE9 mutants (S438P, V176I) that have normal protein expression and endosomal localization but have lost ion transport activity and thus fail to alkalinize the endosomal lumen (Fig. 6A) (24). We validated localization of wild type and mutant NHE9 to the recycling endosome by co-localization with transferrin in GBM 276 (Fig. S3A). Only wild type NHE9, but not S438P and V176I mutants was able to increase Oct4 and Nanog transcript (Fig. 6B). This finding was confirmed by Western blotting: mutants S438P and V176I failed to phenocopy wild type NHE9 in stabilizing IGF-1R and failed to elevate Oct4 and Nanog (Fig. 6C). These observations point to the importance of endosomal Na^+^/H^+^ exchange activity in tumor initiation.

**Figure 6:**
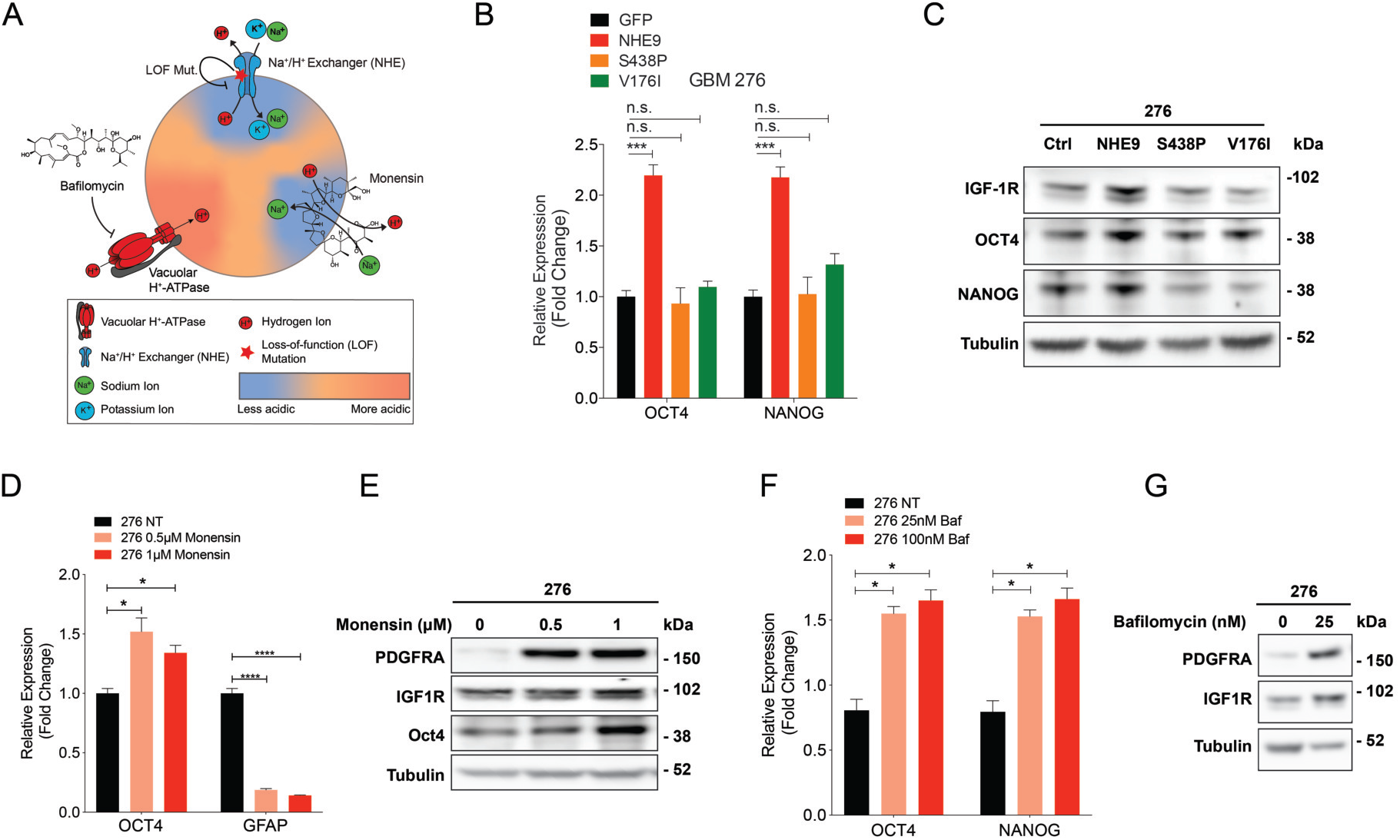
Endosomal pH regulates stemness in glioblastoma cells. (A) Schematic of the three approaches used to query the role of endosomal pH. Loss of function (LOF) mutations in the Na^+^/H^+^ exchanger prevent endosomal alkalinization. Acidification by Vacuolar H^+^-ATPase is blocked by bafilomycin. The ionophore monensin mimics Na^+^/H^+^ exchange activity of NHE9. (B) qPCR analysis of Oct4 and Nanog transcripts in GBM 276 expressing vector (GFP), or the following GFP-tagged constructs: NHE9, S438P, or V176I. (C) Western blot of GBM 276 lysates expressing GFP (Ctrl) or the NHE9 constructs indicated, for IGF-1R, Oct4, and Nanog. (D) qPCR for Oct4 and Nanog transcripts in GBM 276 with no-treatment, 0.5 μM monensin, or 1 μM monensin for 16 h. (E) Western blot of PDGFRα, IGF-1R, and Oct4 in GBM 276 treated with monensin at the concentrations shown for 16 h. (F) qPCR analysis for Oct4 and Nanog transcripts in GBM 276 treated with bafilomycin at the concentrations shown for 6 h. (G) Western blot for PDGFRα and IGF-1R in GBM 276 treated with bafilomycin as shown for 6 h.

The pH of the recycling endosome is precisely tuned by a balance between inward proton pumping through V-ATPase and outward proton leak by NHE9 (Fig.6A). Thus, high levels of NHE9 expression seen in GBM cells corresponds to alkaline endosomes (20). To directly query the role of pH in glioblastoma stemness, we used pharmacological approaches to alkalinize the endosomal lumen and bypass the requirement for NHE9. The ionophore monensin is a Na^+^/H^+^ exchanger mimetic (52, 53) (Fig. 6A). Acute application of monensin to GBM 276 cells caused a dose-dependent elevation of endosomal pH as expected (Fig. S3B- C). Here we show that monensin increased Oct4 transcript relative to vehicle control and caused a reciprocal decrease in GFAP levels (Fig. 6D). Furthermore, monensin also stabilized IGF-1R and PDGFRα, and increased Oct4 protein as seen by Western blotting (Fig. 6E). Bafilomycin inhibits the H^+^ pumping V-ATPase to prevent acidification of endo-lysosomal compartments (Fig. 6A). We show that bafilomycin treatment of GBM 276 increased Oct4 and Nanog transcript levels (Fig. 6F) and stabilized PDGFRα and IGF-1R (Fig. 6G). Taken together, these findings reveal that alkalinization of endosomal pH is the underlying mechanism for NHE9-mediated stemness in glioblastoma.

### NHE9 induces self-renewal capacity in vitro and tumor formation in vivo

The functional consequence of increased stemness is typically quantified *in vitro* using tumorsphere formation as a means to assess self-renewal capacity of cells. As seen in the fluorescence images, tumorspheres formed from GBM 276 cells transfected with NHE9-GFP were larger, consistent with our previous report that NHE9 increases tumor cell proliferation (20), and also more numerous than GFP vector-transfected control (Fig. 7A). Furthermore, the number of spheres per uncoated well correlated with seed density and NHE9 expression: for the same seed density, significantly more spheres were obtained from NHE9-high GBM 612 compared to NHE9-low GBM 276 (Fig. 7B). Tumorsphere formation capacity decreased after NHE9 knockdown in GBM 612 and increased after transgene NHE9 expression in GBM 276 (Fig. 7 C-D). A quantitative estimate of stem cell frequency was determined by extreme limiting dilution assay (ELDA)(54). More progenitor cells capable of seeding tumorspheres were detected in GBM 612, relative to GBM 276 (Fig. 7E). Progenitor cell frequency decreased with NHE9 KD (Fig. 7F) and increased with transgene NHE9 expression (Fig. 7G). Furthermore, unlike wild type NHE9, S438P and V176I mutants do not increase self-renewal capacity (Fig. 7H), and confirm the findings from Figure 6 on the importance of Na^+^/H^+^ exchange activity in conferring stemness phenotypes.

**Figure 7:**
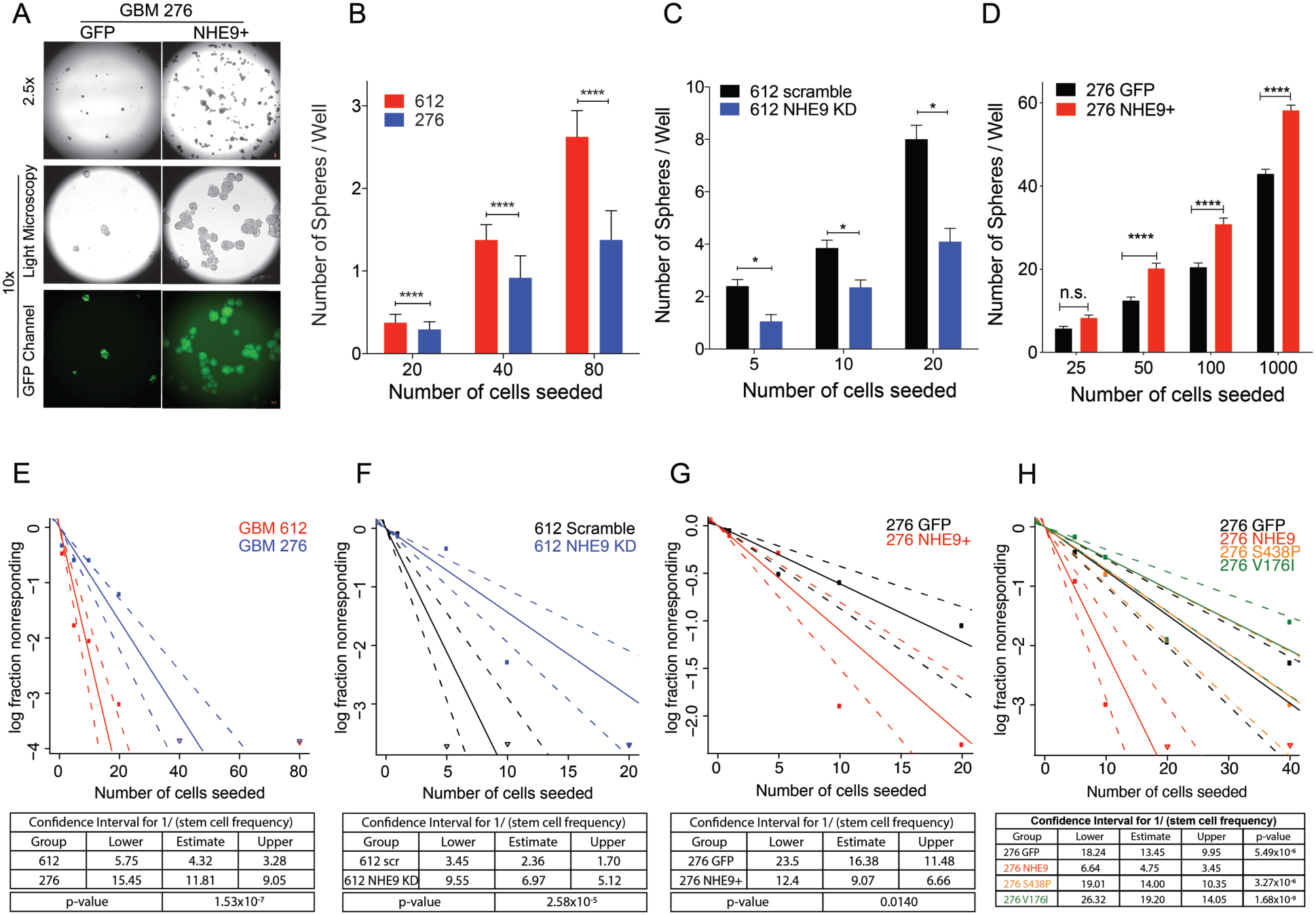
NHE9 increases self-renewal in glioblastoma cells. (A) Light microscopy and fluorescence images of tumorspheres formed from GBM 276 GFP or NHE9-GFP cells in suspension in serum-free media on a non-coated petri dish after two weeks of culturing (Scale bars, 50 μm). (B-D) Number of tumorspheres per well in a 96 well plate at various seeding dilutions as indicated in cell numbers for (B) GBM 612 or GBM 276 (C) GBM 612 scramble or NHE9 KD (D) GBM 276 GFP or NHE9-GFP. (E-H) Extreme limiting dilution assay for estimated stem cell frequency and confidence interval for (E) GBM 612 or GBM 276, (F) GBM 612 scramble or NHE9KD, (G) GBM 276 GFP or NHE9-GFP, and (H) GBM 276 GFP, NHE9-GFP, NHE9-GFP (S438P), or NHE9-GFP (V176I).

Previously, we reported that NHE9 expression is elevated in most GBM subtypes, with highest levels in mesenchymal tumors (20). A small sample of glioblastoma tumor stem cells (n=22) relative to normal neural stem cells (n=3) showed a 5-fold elevation of NHE9 transcript (20). Expanded analysis using cBioPortal shows elevated NHE9 expression in glioblastoma, in comparison to non-malignant brain samples (Fig. 8A). Statistical analysis (ANOVA) of NHE9 expression in 24 immunohistochemistry slides revealed significant variability between normal, low-grade glioma (LGG), and high-grade glioma (HGG) samples (p = 0.0185) as shown by the representative images in Fig. 8B. In addition, student t-test analysis of NHE9 staining between LGG and HGG patients showed significant difference (p = 0.0272).

**Figure 8:**
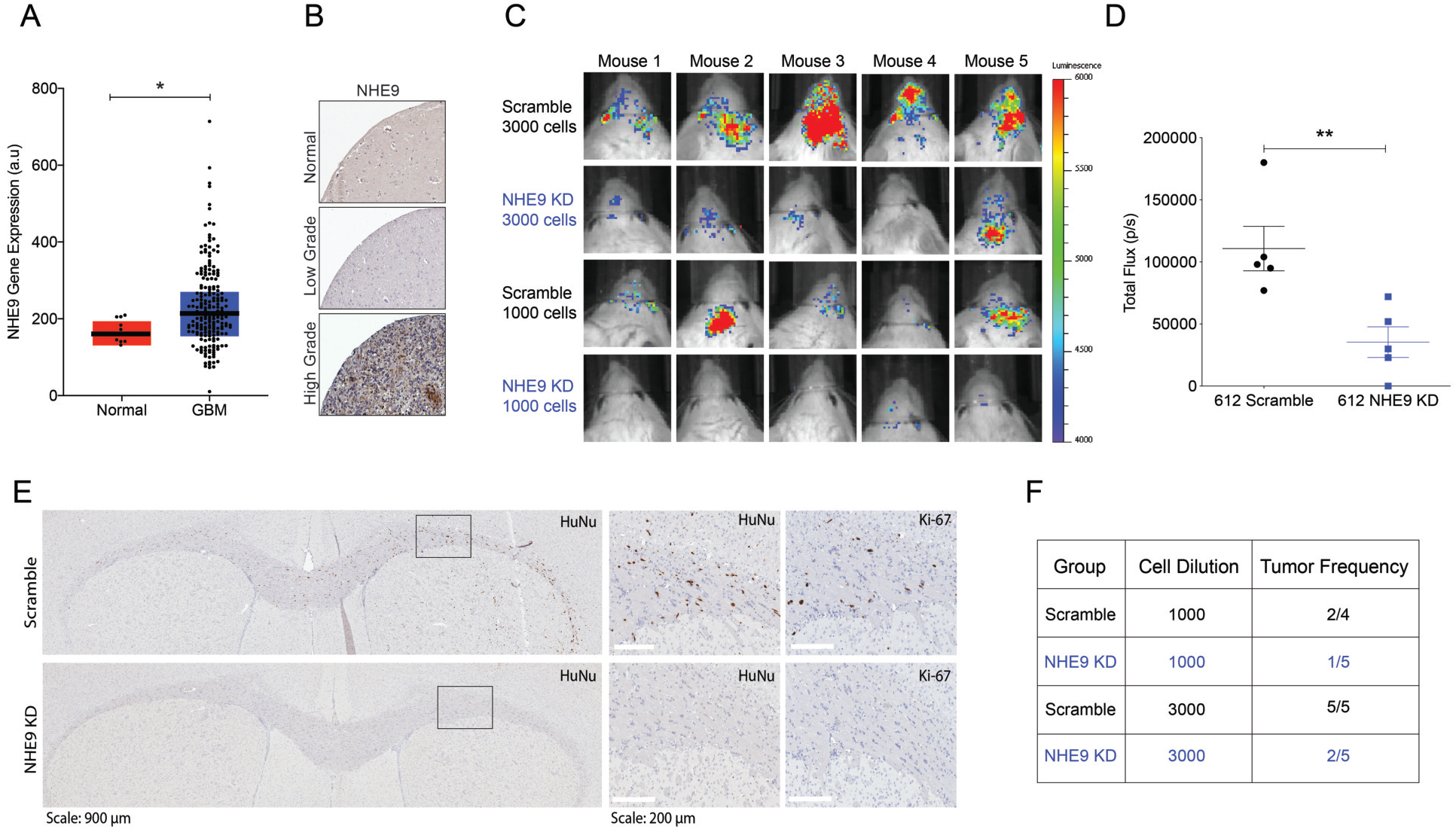
NHE9 increases tumor initiation *in vivo*. (A) Comparison of NHE9 (*SLC9A9*) mRNA expression between GBM tumor and normal samples from cBioPortal database (\**p*-value: 0.0423; Student’s *t*-test). (B) Representative immunohistochemistry staining for NHE9 protein from normal, low-grade glioma, and high-grade glioma, obtained from the Human Protein Atlas. (C) Bioluminescence signal from mice with orthotopic brain xenograft of luciferase-expressing 612 GBM tumor cells with or without NHE9-shRNA seeded with 3000 or 1000 cells. (D) Quantification of total bioluminescence flux from mice with 3000 cells (\*\**p*-value: 0.0084; Student’s *t*-test). (E) Representative immunostaining images for Human Nuclei (HuNu) and Ki-67 from brain tissues injected with control or NHE9 KD cells with 3000 cells. (F) Tumor frequency in mice injected with GBM 612 control or NHE9 KD cells with 1000 and 3000 cells.

To determine the contribution of NHE9 to tumor initiation *in vivo,* we performed orthotopic transplant of luciferase-expressing GBM 612 transfected with scramble RNA (control) or shNHE9 (knockdown, KD) at low seed counts (1000 and 3000 cells). Fig. 8C shows bioluminescence images of mouse brains 10 weeks post-xenograft. Quantitative luminescence analysis revealed significantly decreased light flux in mouse brains injected with NHE9 KD cells compared to control pointing to fewer tumor cells (*p* = 0.0084; 3000 cells) (Fig. 8D). Subsequently, brains were sectioned and stained for human nuclei (HuNu) and the cell proliferation marker Ki67 to evaluate tumor formation. Representative immunohistochemistry images are shown in Fig. 8E. For control mice, immune-positive staining was found for both HuNu and Ki67 markers across the corpus callosum and dispersed throughout the brain parenchyma. Conversely, mice injected with NHE9 KD cells showed little or no staining. Total body weight of all mice was similar and remained mostly unchanged over 11 weeks (Fig. S4A). After investigating all stained brain sections (Fig. S4B-C) for viable, dividing human GBM cells, we confirmed decreased tumor frequency with NHE9 KD (Fig. 8F). Quantitative analysis of these data revealed statistically significant difference in tumor formation (*p* = 0.0185). These findings demonstrate that NHE9 not only elevates expression of genes associated with stemness and pluripotency, but also self-renewal capacity at a functional level, measured by tumorsphere formation *in vitro* and tumor initiation *in vivo*.

## DISCUSSION

The recycling endosome is a critical hub for oncogenic receptor trafficking. There is accumulating evidence that dysregulation of endosomal trafficking in cancer contributes to the progression of malignancy. For example, quantitative proteomics of primary brain tumor samples revealed that the endocytic machinery is commonly down regulated in brain malignancies regardless of tumor grade and type (55). Intriguingly, loss of the RNA-binding protein QKI was shown to attenuate degradation of oncogenic growth factor receptors by down regulating biogenesis of endo-lysosomal compartments, leading to maintenance of self-renewal in glioma stem cells even in a growth factor-deficient environment (56). Here, we show that gain of NHE9 function and hypo-acidification of the recycling endosome lumen increases recycling of oncogenic growth factor receptors and promotes their escape from degradation, resulting in tumor stemness phenotypes. In another study, a suboptimal environment of iron-deficiency in tumors was found to increase NHE9-mediated transferrin receptor surface recycling in a model of the blood-brain barrier (57). Thus, brain tumor cells adopt multiple mechanisms of altered endosomal trafficking to sustain oncogenic receptor density at the cell surface and scavenge scarce nutrients from the tumor microenvironment. Despite their recognized role in cell-environment signaling, recycling endosomes have not been therapeutically exploited, due to a lack of a specific and selective target.

We describe a mechanism of endosomal pH dysregulation through an imbalance of proton pump and leak pathways that allows multiple receptors to evade lysosomal degradation, expanding on our previous observation linking NHE9 to epidermal growth factor receptor (EGFR) expression levels (20). By alkalinizing the endosomal lumen in glioblastoma cells, NHE9 functionally recapitulates the role of E5 oncoprotein of human papilloma virus (HPV), the main cause of cervical cancer worldwide. E5 interacts with V-ATPase to inhibit proton pump activity, alkalinize endosomes and drive tumorigenesis (58–61). However, tumorigenic alterations in pH appear to be compartment-specific: in a model of breast cancer, decreased expression of NHE6 correlated with hyper-acidification of early endosomes resulting in drug sequestration and chemotherapeutic resistance (21). On the other hand, elevated expression of the *trans*-Golgi network isoform NHE7 in a pancreatic cancer model also resulted in hyperacidification of compartmental lumen and activation of plasma membrane Na^+^/H^+^ exchange (22), mimicking NHE1-mediated regulation of cytoplasmic pH and extracellular acidification common to a wide range of cancer types.

Although hyperactivation of oncogenic receptor tyrosine kinase signaling in glioblastoma tumors has been extensively appreciated (62, 63), efforts in therapeutic targeting of single receptors such as EGFR and PDGFR have largely failed in clinical trials due to receptor-heterogeneity and redundancy. Finding a drug target that inhibits multiple oncogenic receptors may circumvent this problem (64, 65). Combination therapy - targeting multiple RTKs, has been suggested (12) but the plasticity of tumor cells and their stochastic response has prevented this approach from being effective and approved for patients. Furthermore, the existence of a small population of cells in glioblastoma that has stem cell-like characteristics drives tumor recurrence (8). Thus, a druggable target such as NHE9 that controls both stemness and pan-receptor clearance in GBM cells could circumvent shortcomings in current therapy. The use of weak bases like chloroquine or V-ATPase inhibitors that de-acidify endo-lysosomes is being explored as therapeutic adjuvant therapy in GBM (66, 67). Our findings could spur a search for inhibitors that selectively target the NHE9 isoform.

The link between pH and pluripotency is intriguing but not fully explored. Few studies suggest that exposure to extracellular acidity can induce stemness in tumors (14, 68), however the role of organellar pH is unclear. Furthermore, the link between extracellular and endosomal pH, if any, should be investigated in future studies. This work represents a first step in evaluating the role of NHE9 in mediating stemness, and as such, has some limitations. Strictly speaking, stem cells are capable of self-renewal as well as multilineage differentiation therefore a thorough evaluation of molecular and functional lineage and stem cell markers (13, 69) associated with NHE9 in GBM may be helpful. While significant effort has been invested in deciphering the role of extracellular pH in various malignancies, our findings revealing the previously unexplored role of endosomal pH in stemness warrant further study.

## METHODS

### GBM Cell Culture

All glioblastoma tumor samples and subsequent cell line derivations were performed with written consents from patients through Quinones laboratory, observing Institutional Review Board guidelines as previously described. Glioblastoma stem cells with tumorsphere forming capabilities, GBM 276 and GBM 612, were cultured in DMEM/F12 (1:1), 1% HyClone Antibiotic Antimycotic Solution (Thermo Scientific), Gibco™ B-27 serum free Supplement (Thermo Scientific 17504044), 20ng/ml EGF, and 20ng/ml FGF. For ligand treatment experiments, Gibco™ B27 serum free supplement, minus insulin (Thermo Scientific A1895601) was used. For adherent culture, dishes were coated with Laminin (Sigma-Aldrich L2020). Coating was done in DMEM/F12 basal media for at least 2 h or overnight, and for every 1 cm^2^ of surface area, 1 μl of laminin was added.

### Plasmids

Full-length mNHE9 was cloned into FuGW lentiviral vector as previously described. HsNHE9 targeting short hairpin RNA (shRNA) (5’-CCGGCCCTCCATTAAGGAGAGTTTTTCAAGAGAAAACTCTCCTTAATGGAGGTTTTTC-3’) and scramble control (Sigma-Aldrich) (5’-CAACAAGATGAAGAGCACCAA-3’) were cloned into pLKO.1 lentiviral vector with ampicillin and puromycin-resistance genes for bacteria selection and mammalian cell selection respectively. Constitutively active STAT3 (EF.STAT3C.Ubc.GFP, plasmid#24983) was purchased from Addgene.

### Production of lentivirus and cell transduction

pCMV-dR8.9 (8.8μg), pMD2.G (2μg) lentivirus packaging plasmids, and NHE9 plasmids (10 μg) were co-transfected with pLKO.1-shNHE9 in HEK293FT cells with lipofectamine 3000 (ThermoFisher Scientific) according to manufacturer’s protocol. Media was changed the next day, and virus was collected 48h, 72h, and 96h after transfection. The collected media containing virus was filtered through 0.45μm pore filter and mixed with Lenti-X concentrator (Clontech) as indicated by manufacturer’s instruction. Centrifugation was preformed overnight incubation at 4°C at 3300 rpm for 15 mins. The pellet was resuspended in PBS and was either used immediately or frozen at −80°C for later use. For glioblastoma cell transduction, cells were seeded on laminin-coated dishes and infected with lentivirus. In case of shRNA constructs, cells were incubated with lentivirus for 48 h in low volume of media. Subsequently, GBM612 cells were recovered in GBM complete media for 6 h and were selected with puromycin (1μg/ml). GBM 276 cells transduced with FuGW and mNHE9-GFP virus were sorted by fluorescence-activated cell sorting (FACS) for GFP positive cells to be used for subsequent assays and analysis.

### Quantitative PCR analysis

RNA was extracted from cultured cells using the RNeasy Mini kit (#74104, Qiagen) according to the manufacturer’s instructions. Complementary DNA was synthesized using the high-Capacity RNA-to-cDNA kit (#4387406, Applied Biosystems). Quantitative PCR analysis was performed using the 7500 Real-Time PCR system (Applied Biosystems) with Taqman 2x Fast universal PCR Master Mix (#4304437, Applied Biosystems). Taqman gene expression assay probes used in this study are: Human: NHE9 (SLC9A9), Hs00543518; NHE6 (SLC9A6), Hs00234723; Oct4 (POU5F1), Hs00999632; Nanog, Hs04399610; Sox2, Hs01053049; c-Myc, Hs00811069;KLF4, Hs00358836;Olig2, Hs00300164; Nestin (NES), Hs04187831; CD133(PROM1), Hs01009257; GFAP, Hs00909233;TUJ1 (TUBB3), Hs00801390;O4 (FOXO4), Hs00936217; IGF1R, Hs00609566; IR, Hs00961557; PDGFRA, Hs00998018; GAPDH, Hs02786624. The Ct (cycle threshold) values were determined first and subsequently normalized to GAPDH expression (endogenous control) to obtain ΔCt value for each sample. These ΔCt values were compared to the control ΔCt values and acquired ΔΔCt values. Fold changes were then calculated using the equation: expression fold change = 2 ^− ΔΔCt^. Each experiment was performed at least three times independently.

### Immunoblotting

Cells were lysed using Pierce™ RIPA buffer (#89900, Thermo Scientific) supplemented with Protease/phosphatase Inhibitor cocktail (100x) (5872S, Cell Signaling Technology). Cells were sonicated for 15-20 seconds each sample and rotated for 15 min at 14,000 rpm at 4°C. Protein concentration was determined by the BCA assay. For each sample, 15-30 μg of protein was mixed with 10x reducing buffer and 4x loading buffer. The samples were heated for 10 min at 70 °C and separated on 10 or 12 well NuPage™ 4-12% Bis-Tris Gel (NP0322BOX or NP0321BOX) under reducing conditions, and then transferred onto activated PVDF membranes. Ponceau staining was utilized to confirm protein transfer. The membranes were blocked with either 5% milk or 5% BSA overnight at 4°C. The membranes were subsequently incubated with primary antibodies overnight at 4°C, followed by 1h incubation with HRP-conjugated secondary antibodies. SuperSignal™ West Pico PLUS Luminol/Enhancer solution and Stable Peroxide solution were mixed with 1:1 ratio and were introduced to the membranes for HRP activation. For imaging, digital Amersham Imager 600 system was used. Densitometric quantification and image processing was performed with ImageJ software or FIJI. Human Phospho-RTK Array Kit (ARY001B, R&D Systems) was used according to the manufacturer’s instructions and the blots were imaged and quantified in the same way as western blots.

### Immunofluorescence

PHEM buffer was prepared (60mM PIPES, 25mM Hepes, 10mM EGTA, and 2mM MgCl_2_ at pH 6.8). Cells were cultured on glass coverslips and were rinsed with PBS and pre-extracted with 1X PHEM buffer, 8% sucrose and 0.025% saponin. Cells were fixed with 4% paraformaldehyde in 1x PBS for 30 min and were washed with PBS 3 times for 5 min each. After blocking in 1% BSA in PBS with 0.025% saponin, cells were incubated with primary antibody in 1% BSA in 1x PBS with 0.025% saponin overnight at 4°C. Cells were rinsed with 0.2% BSA in 1x PBS 3 times for 10 min each and were then incubated with a fluorescent secondary antibody (1:1000) in 1% BSA in 1x PBS with 0.025% saponin for 30 min at RT. Coverslips were washed 3 times for 5 min each with 0.2% BSA in 1xPBS and mounted onto slides with ProLong(R) Gold Antifade with DAPI Molecular Probes (8961S, Cell Signaling Technology). Images were taken with Zeiss LSM780-FCS Single point, laser scanning confocal microscope and the obtained images were analyzed using ImageJ.

### Endosomal pH measurement

Endosomal pH measurement was measured using pH sensitive FITC (50 μg/ml), normalized for uptake with pH insensitive Alexa Fluor 633 (50 μg/ml) (Invitrogen). tagged to transferrin (100 μg/ml) as described previously (20). Where indicated, treatment of GBM 276 cells with monensin (10 and 50 μM) was for 1 hr. Adherent cells were washed with PBS once, treated with tagged transferrin at 37 °C for 55 min. Transferrin uptake was stopped by chilling the cells on ice. Excess transferrin was removed by washing with ice-cold PBS, whereas cell surface bound transferrin was removed by an acid wash in PBS at pH 5.0 followed by a wash with PBS at pH 7.4. The fluorescence intensity for both dyes was measured for at least 5,000 cells by flow cytometry using the FACSAria (BD Biosciences) instrument and the average intensity of the cell population was recorded. A pH calibration curve was generated in buffers at different pH (4.5, 5.5, and 6.5) in the presence of K^+^/H^+^ ionophores, nigericin (10 μM) and valinomycin (10 μM) by following manufacturer’s protocol from the Intracellular pH Calibration Buffer Kit (Invitrogen).

### Extreme limiting dilution assay (ELDA)

The cells were enzymatically dissociated with Accutase and were filtered into 5ml polystyrene round-bottom tube with cell strainer cap (#352235, Falcon). In order to increase the accuracy of the number of cells seeded, the FACS Aria sorter at the Ross Flow Cytometry Core at the Johns Hopkins was utilized to seed indicated number of cells in each well of uncoated 96 well plate. The cells were cultured in sphere form for two weeks, and the number of spheres was determined for each well. Analyses of self-renewal ability and stem cell frequency were performed using the algorithm on the following website: http://bioinf.wehi.edu.au/software/elda/index.html.

### Intracranial implantation of GBM cells

In vivo methods were carried out in accordance to the JHU Institutional Handbook on the Use of Experimental Animals. Intracranial implantation of GBM612 cells (transduced with control and NHEKD vectors) was carried out using two different dilutions of 1000 cells (n=9) and 3000 cells (n=10). Orthotopic implantation of cells was done in 6-week-old male athymic nude mice. Animals were anesthetized with ketamine HCl (80mg/kg) and xylazine HCl (8mg/kg) via IP injection. Mice were then positioned in a stereotactic frame (KOPF model 2006 small animal stereotaxic instrument with digital display console) with a surgical microscope (Zeiss surgical microscope MPMI-1 FC). A linear midline incision of the scalp was made to expose the calvarium. Different dilutions of GBM612 cells were stereotactically injected into the striatum *(X: 1.5mm, Y: 1.34mm, Z: −3.5mm from Bregma)*. After cell injection, the needle was carefully retracted, and the skin was closed with surgical glue. Mice were placed on a heating pad to speed recovery. Tumor formation and growth was confirmed by weekly bioluminescence (BLI). After 11 weeks of surgical implantation of tumors, mice were intracardially perfused with 4% paraformaldehyde. Tumor formation was determined by immunohistochemistry.

### Immunohistochemistry

Mouse brain tissue was fixed in 4% PFA and embedded in paraffin. Formalin-fixed paraffin embedded sections (10 μm thick) were deparaffinized and rehydrated, followed by antigen recovery. 3% hydrogen peroxide was used to block endogenous peroxidase activity, followed by Rodent Block M (Biocare) after which slides were washed in phosphate-buffered saline and stained against Human Nuclei (MAB4190, 1:250, Millipore) and Human specific Ki67 (MAB4383, 1:500, Millipore), followed by secondary antibody Envision+ anti-mouse labeled-polymer (Dako/Agilent). Slides were counterstained using Gill’s I hematoxylin (Richard-Allen). Stained slides were imaged using Aperio AT2 scanner and analyzed by Aperio ImageScope software.

### Statistical analysis

All experiments were conducted at least in triplicates to meet the standard for statistical power. For quantitative PCR analysis, ΔΔCt values in triplicate were calculated by Applied Biosystems 7500 analysis program. Those values were normalized in comparison to the control, which then was analyzed by using two tailed Student’s t-test. For western blotting analysis and flow cytometry, normalized densitometry and mean fluorescent values, respectively, were analyzed with two tailed Student t-test. Statistical significance was assumed when the p-value was lower than 0.05. (non-significant: p-value > 0.5, *: p-value <0.05, **: p-value <0.01, **: p-value <0.001, and ****: p-value <0.0001).

## Acknowledgements

This work was supported by the following grants: NIH R01DK108304 and US-Israel Binational Science Foundation grant (RR) and Mayo Clinic Investigator Award (AQH), State of Florida Cancer Research Award (AQH), NIH R01 NS070024 (AQH). The following fellowship support is also acknowledged: Ruth L. Kirschstein Individual National Research Service Award F31CA220967 (MJK), American Association of Cancer Research-Astra Zeneca Breast Cancer Research Fellowship (MRM) and HHMI Gilliam Fellowship for Advanced Study (AXM). We thank Dr. Shuli Xia and Dr. Junhua Yang for helpful discussions, and Dr. Xiaoling Zhang and Dixie Hoyle of Ross Flow Cytometry Core at the Johns Hopkins School of Medicine for technical support.

## Notes

### Competing Interest Statement

The authors have declared no competing interest.

