## Supplemental Figures for "The Endosomal pH Regulator NHE9 is a Driver of Stemness in Glioblastoma"

**Figures S1-S4**

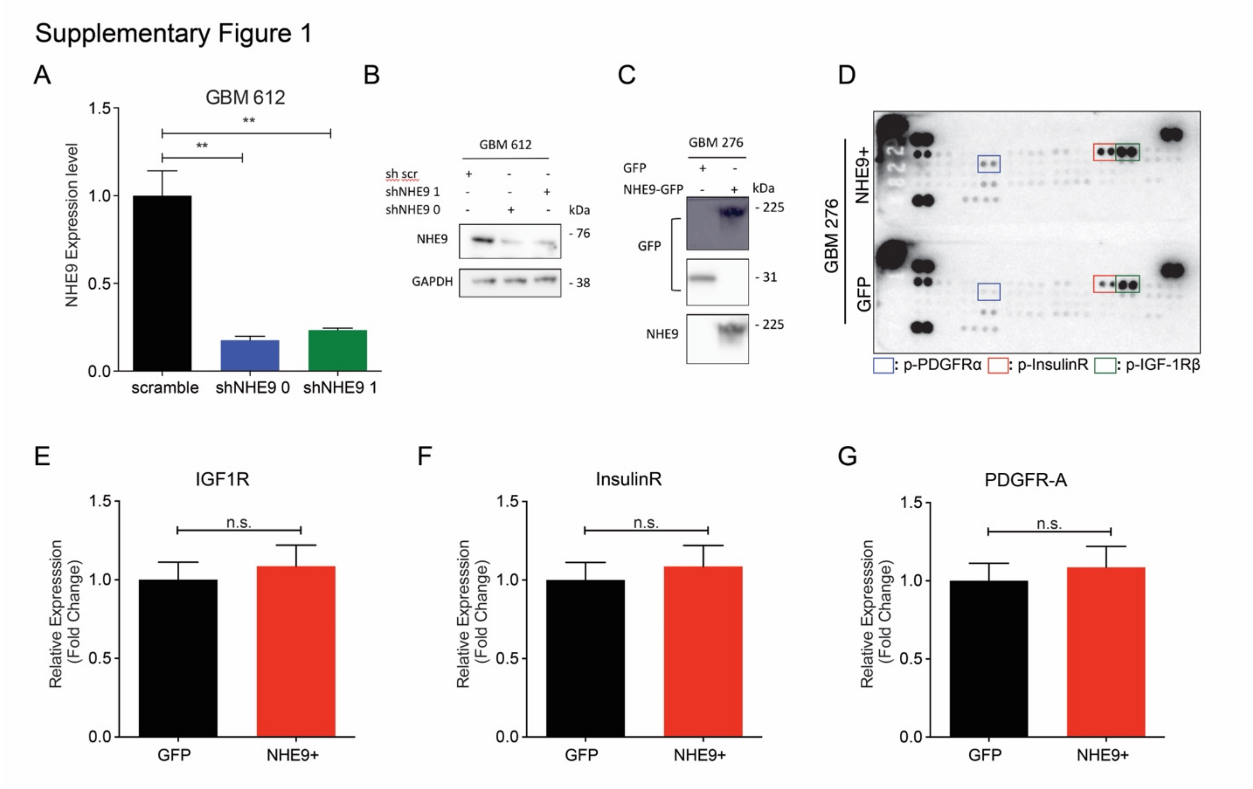


**Figure S1**: **Effect of NHE9 expression in GBM on receptor tyrosine kinases**

Relates to Figure 1.

(A) qPCR for NHE9 transcript in GBM 612 cells following treatment with two targeting shRNA constructs, relative to scramble control. ***p*-value: 0.0046 - Student's *t*-test; ***p*-value: 0.0057, Student's *t*-test). (B) Western blot of NHE9 of samples from A. (C) Western blot for GFP (top two panels) and NHE9 (bottom panel) in GBM 276 cells transfected with GFP or NHE9-GFP. (D) phosphorylated RTK dot blot overlaid with cell lysate from GBM 276 transfected with GFP or NHE9-GFP. Boxed samples show p-RTKs excerpted in Fig. 1C, identified as shown by colored boxes, and chosen for study. (E-G) qPCR analysis for transcript levels of (E) IGF-1Rβ, (F) InsulinR, and (G) PDGFRα.

Supplementary Figure 2

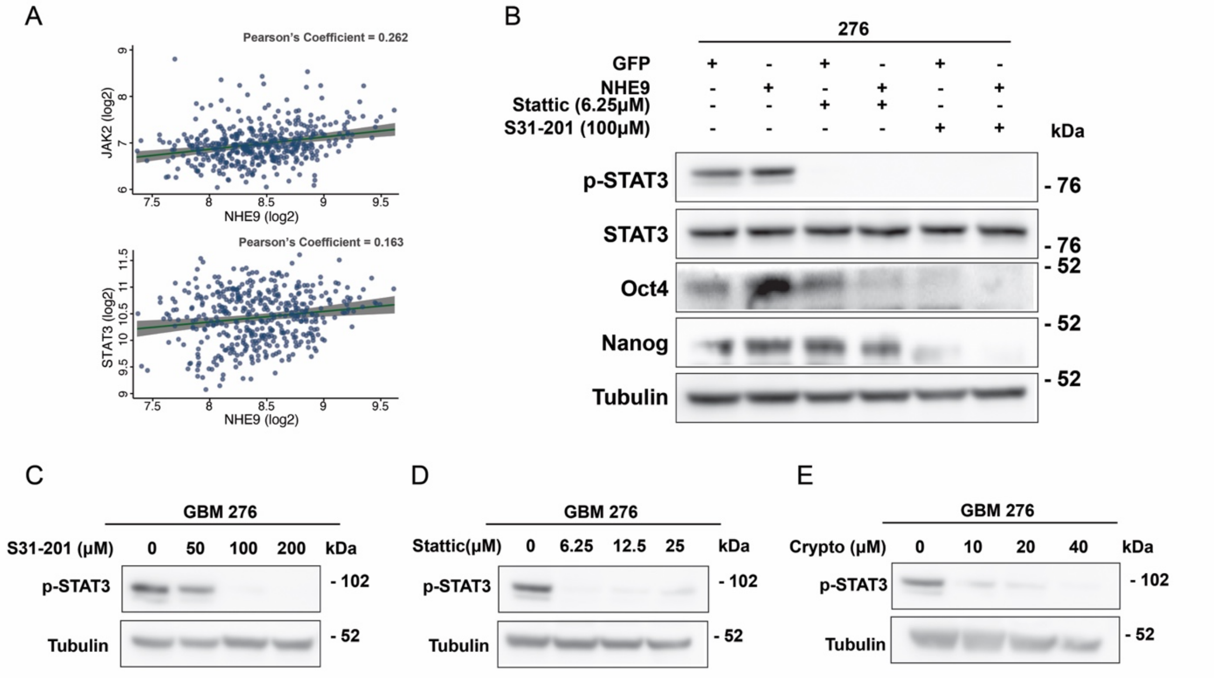


**Figure S2: Analysis of STAT3 and JAK2 role in GBM**

Relates to Figure 3.

(A) Correlation analysis between NHE9 and JAK2 (top) and STAT3 (bottom) transcript levels in GBM patients obtained from Rembrandt dataset on Betastasis (JAK2/NHE9 transcript Pearson's coefficient: 0.262; STAT3/NHE9 transcript Pearson's coefficient: 0.163). (B) Western blot for p-STAT3, STAT3, Oct4, and Nanog proteins in GBM 276 GFP and NHE9-GFP, with STAT3 inhibitors, Cryptotanshinone or Stattic as indicated. (C-E) Western blot for p-STAT3 in GBM 276 with STAT3 inhibitors at increasing concentrations: (C) 0, 50, 100, and 200 µM of S31-201; (D) 0, 6.25, 12.5, and 25 µM of Stattic; and (E) 0, 10, 20, and 40 µM of Crytptanshinone.

Supplementary Figure 3

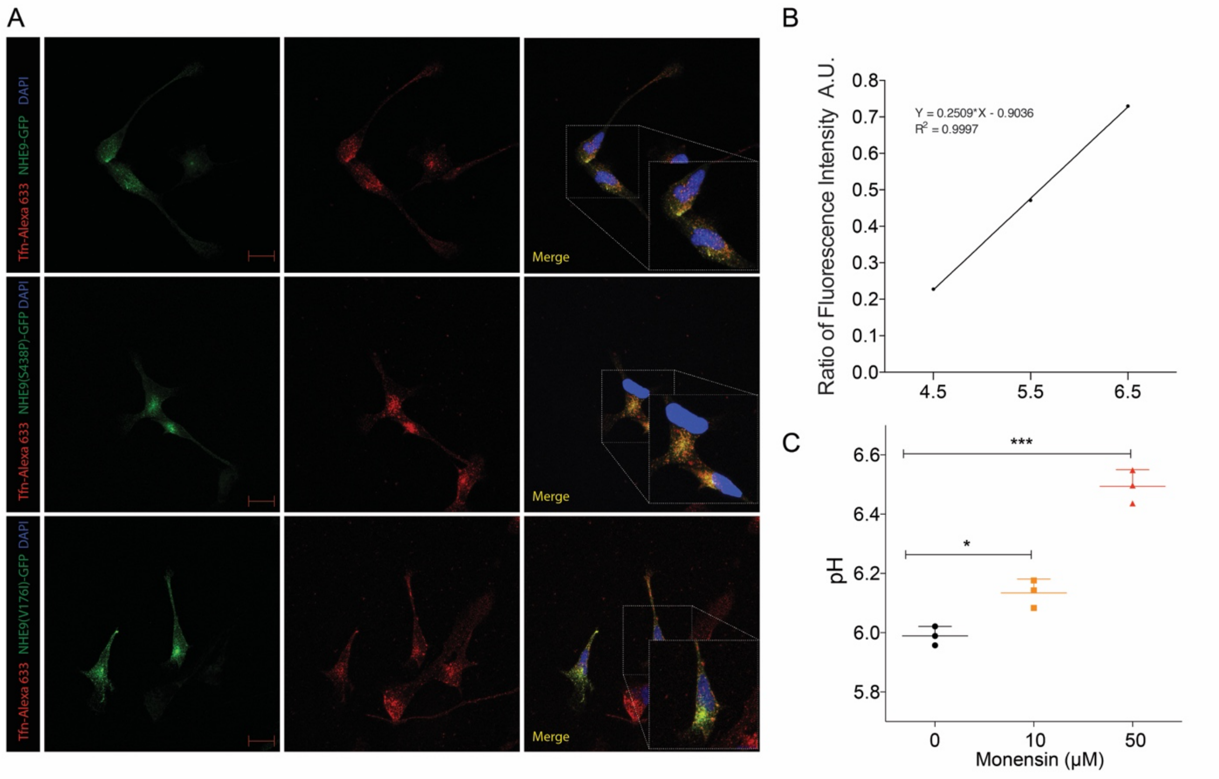


**Figure S3: Role of endosomal pH in GBM**

Relates to Figure 6.

(A) Immunofluorescence images of GBM 276 cells expressing NHE9-GFP, NHE9 (S438P)-GFP, or NHE9 (V176I)-GFP (Green) after 1hr treatment with Transferrin conjugated with Alexa 633 (Red). Nuclei are stained with DAPI (blue). Inset shows merged images; scale bars, 10 µm. (B) Calibration of fluorescence ratio intensity of transferrin tagged FITC and Alexa Fluor 633 in GBM 276 cells using buffers of indicated pH, as described in Methods. (C) Recycling endosome pH in GBM 276 cells treated with the indicated concentrations of monensin for 1 hour, as described in Methods.

Supplementary Figure 4

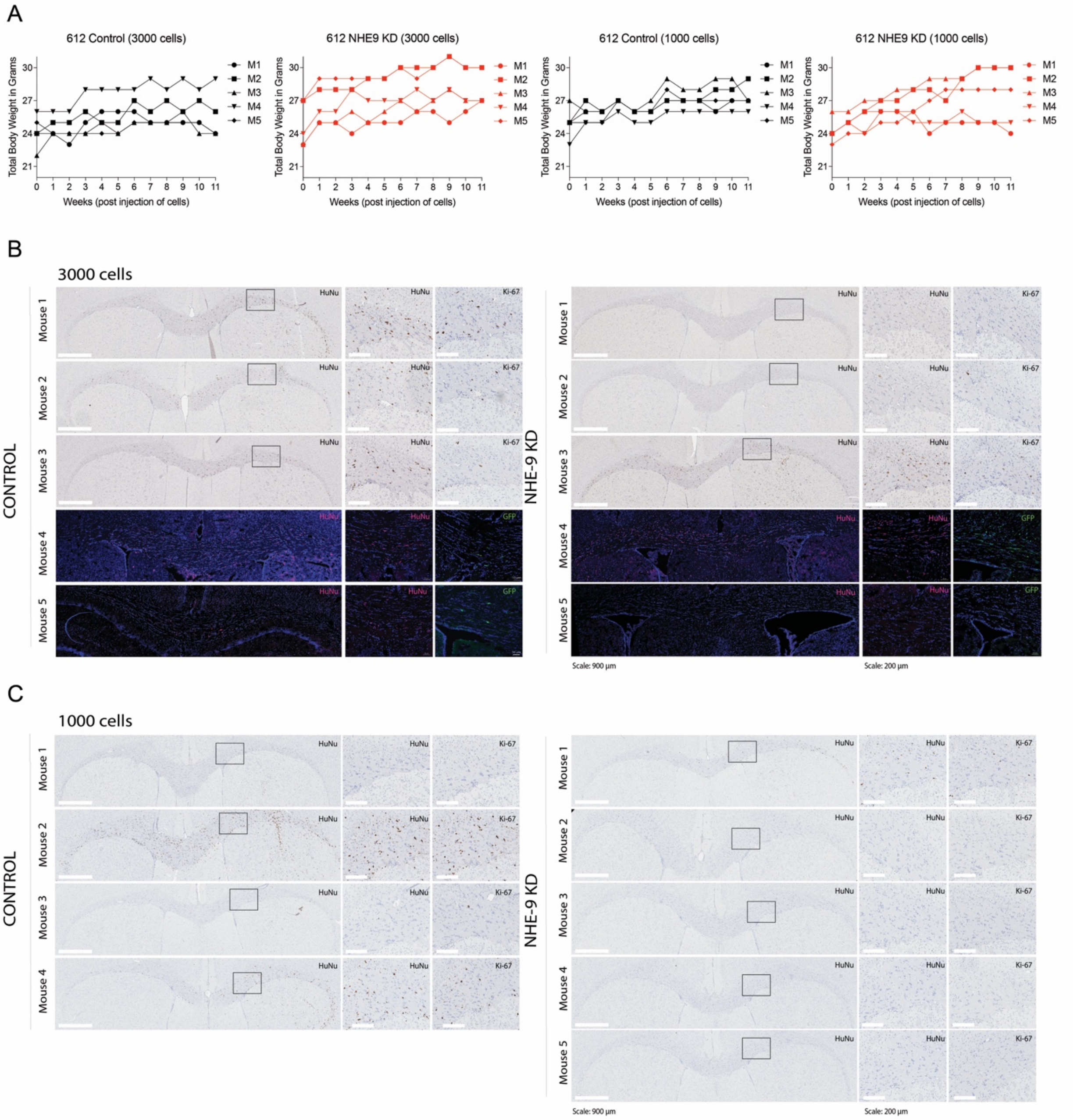


**Figure S4: Effect of NHE9 on tumor initiation in mice**

Relates to Figure 8.

(A) Total body weight of nude athymic mice injected with GBM 612 control or NHE9 KD cells (3000 or 1000), measured in grams over 11 weeks. (B-C) Immunohistochemistry (IHC) and immunofluorescence (IF) images of sectioned brains from mice injected with GBM 612 cells transduced by lentivirus packaged with empty vector or shNHE9 construct at either 3000 cell dilution (A) or 1000 cell dilution (B). (Scales for IHC, 900 µm or 200 µm; for IF, 50 µm).
